# Best-compromise nutritional menus for childhood obesity

**DOI:** 10.1101/618108

**Authors:** Paul Bello, Pedro Gallardo, Lorena Pradenas, Jacques A. Ferland, Victor Parada

## Abstract

Childhood obesity is an undeniable reality and has shown a rapid growth in many countries. Obesity at an early age not only increases the risks of chronic diseases but also produces a problem for the whole healthcare system. One way to alleviate this problem is to provide each patient with an appropriate menu that can be defined with a mathematical model. Existing mathematical models only partially address the objective and constraints of childhood obesity; therefore, the solutions provided are insufficient for health specialists to prepare nutritional menus for individual patients. This manuscript proposes a multiobjective mathematical programming model to aid healthy nutritional menu planning to prevent childhood obesity. This model enables a plan for combinations and amounts of food across different schedules and daily meals. This approach minimizes the major risk factors of childhood obesity (i.e., glycemic load and cholesterol intake). In addition, it considers the minimization of nutritional mismatch and total cost. The model is solved using a deterministic method and two metaheuristic methods. To complete this numerical study, test instances associated with children aged 4-18 years old were created. The quality of the solutions generated using the three methods was similar, but the metaheuristic methods provided solutions in less computational time. The numerical results indicate proper guidelines for personalized plans for individual children.

## 1. Introduction

Childhood obesity has shown a rapid growth in many countries, but this growth can be partially mitigated via optimization mathematical models. This noncommunicable disease is a major concern in public health because being obese at an early age increases the risks of cardiovascular, pulmonary, metabolic, gastrointestinal, skeletal, psychological and other diseases in adulthood [1][2][3]. Also, evidence reveals a positive correlation between obesity/being overweight in childhood and these conditions in adulthood [4]. Therefore, interventions during childhood have great potential because healthy eating habits can be developed during this stage [5]. To address this problem, a health professional must specify the combination and amount of food that the patient should consume at different meal times during the day to ensure the appropriate intake of the nutrients of interest during the planning period. These facts introduce a particular type of operational research problem called the Nutritional-Menu Planning Problem (NMPP). The NMPP is an NP-Hard problem [6] and, in practical terms, the usual method to solve it consists of manually constructing menus through a trial-and-error process that is extremely inefficient and does not guarantee an appropriate menu for each patient.

NMPP variants approached by mathematical models have different objective functions. Stigler [7] and Dantzig [8] were the first to propose the goal of minimizing the total cost of the diet problem. Bas [9] studied the minimization of a risk factor for patients with high glycemic load and metabolic diseases. Orešković, Kĳusurić, and Šatalić [10] maximized the palatability of a menu based on patient preferences by assigning a weight to the objective function in the specific case of vegetarian menus. Masset et al. [11] and Okubo et al. [12] minimized the difference between the quantities currently ingested and the recommended amount while satisfying nutritional requirements. Complementary diets for 6- to 24-month-olds [13] and the planning of nutritional menus at a school in Southeast Asia for 13- to 18-year-olds were studied considering total cost minimization [14]. In some situations, cost minimization alone is insufficient to obtain the proper diet. Other objectives are also relevant, which leads to the multiobjective NMPP that we denote as MO-NMPP.

Several MO-NMPP studies have been conducted. A multiobjective model represents the real problem addressed by the NMPP more completely. Koroušić [6] [15] addressed both economic and aesthetic aspects when generating food menus. The multiobjective model optimizes cost, functionality, seasonality, and other aspects such as flavor, consistency, color, temperature, shape and method of preparation. Donati et al. [16] presented a multiobjective model to generate diets at the lowest cost while minimizing the environmental effect of its production, measured as equivalent carbon dioxide emissions and land and water use. Van Mierlo, Rohmer and Gerdessen [17] studied a similar situation that minimized fossil fuel depletion instead of cost minimization. They found that the existing models for MO-NMPP are focused on general issues that are valid for an obese individual. However, childhood obesity treatments must consider the child’s development. Thus, it is not advisable to provide notably restrictive diets in terms of calories because children are developing. Besides, the recommended menu must encourage the development of healthy eating habits while minimizing exposure to risk factors such as energy-dense, high-fat, high-sugar and high-salt foods. Furthermore, an appropriate glycemic load and an average daily cholesterol intake are necessary. Moreover, the minimum nutritional mismatch between the nutritional contributions provided by the menu and the amount recommended by specialized organizations is an essential condition that the best compromise solution must satisfy. By including all of these components in the original multiobjective problem, we introduce the Multiobjective Nutritional-Menu Planning Problem for Childhood Obesity (MO-NMPP-CHO).

This paper proposes an approach for the MO-NMPP-CHO that considers the minimization of the main risk factors for the development of chronic childhood obesity. The concept of nutritional mismatch is considered, which slightly relaxes the constraints. Also, to avoid limiting the applicability of the menus to sectors with less economic income, the classic objective of minimizing the average daily cost of the menu was considered, which adds to the nutritional constraints suggested by specialized organizations. With the help of a specialist, we created a set of numerical instances that were solved using a deterministic method and two metaheuristic methods.

The remainder of this paper is organized as follows. Section 2 introduces the proposed multiobjective mathematical programming model that seeks to control and prevent childhood obesity. Next, different techniques to solve the problem are summarized in section 3. Section 4 includes the numerical experimentation and discussion. Finally, section 5 presents the main conclusions of this study.

## 2. A multiobjective approach for MO-NMPP-CHO

This section presents the model for the MO-NMPP-CHO. The proposed approach minimizes the main risk factors for the development of the chronic diseases associated with childhood obesity, nutritional mismatch and the average daily cost of the generated menus. The definitions of the parameters and variables present in the model are summarized in Table 1.

**Table 1:**
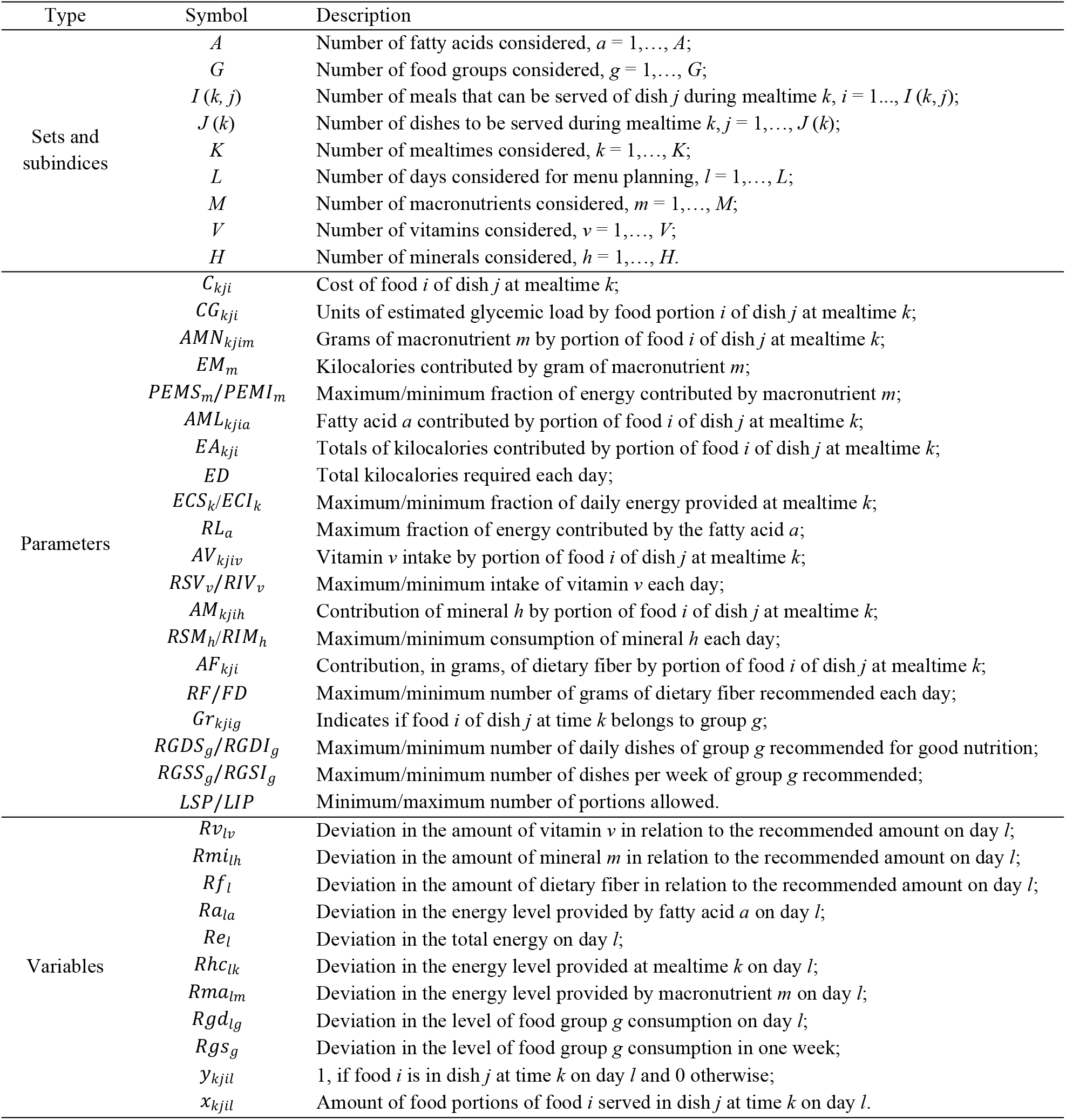
Parameters and decision variables for the MO-NMPP-CHO model

The model that enables the generation of food plans for children to reduce the risk of childhood obesity is presented in equations (1)–(19).

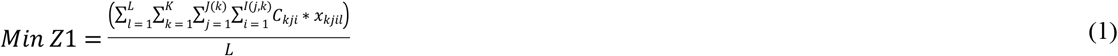

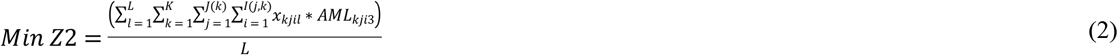

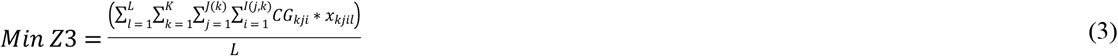

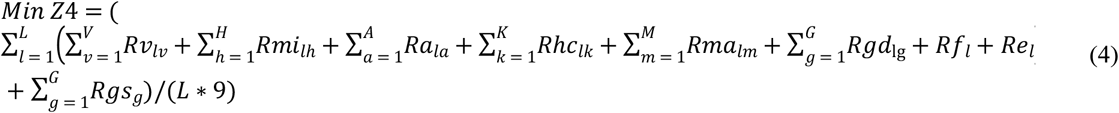

Subject to,

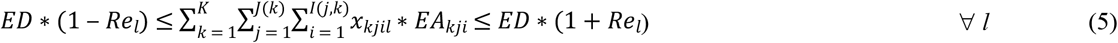

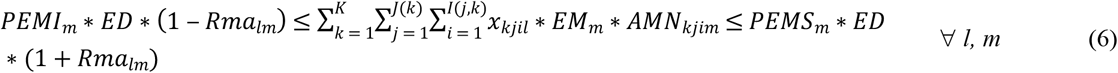

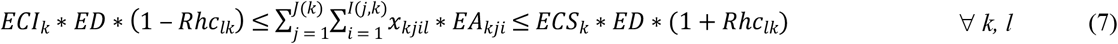

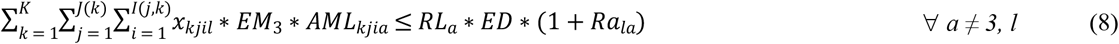

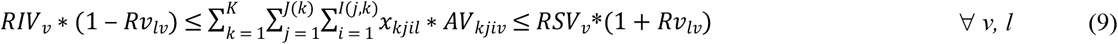

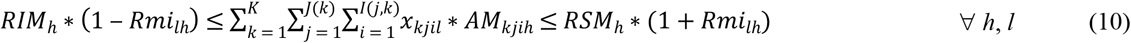

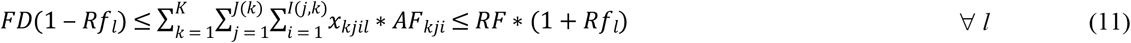

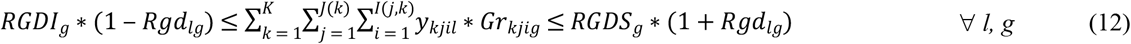

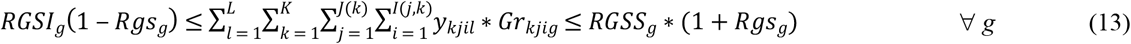

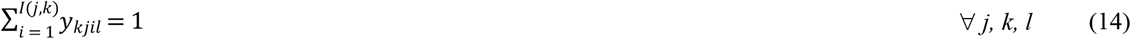

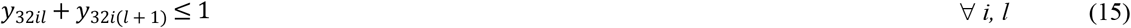

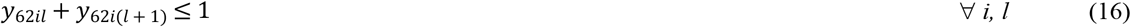

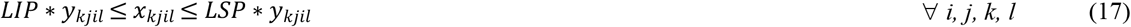

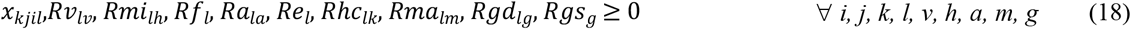

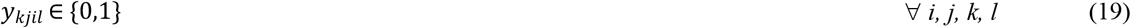

The first four equations, (1) to (4), correspond to the objective functions. The first objective function (1) minimizes the average daily cost of the food plan [7]. The second objective function (2) minimizes the average daily cholesterol intake to reduce the negative effects of fat consumption. The third objective function (3), which was proposed by Bas [9], minimizes the average daily glycemic load of the menu. The glycemic load (GL) corresponds to the glycemic index (*GI*), adjusted by a specific amount of carbohydrates (*GL* = carbohydrates x *GI*/100). This concept is of interest because the consumption of low-glycemic-index foods reduces the risk of diseases associated with hyperinsulinemia (excess insulin in the blood) such as diabetes mellitus and cardiovascular diseases, while also decreasing the sensation of hunger [18]. Finally, the fourth objective (4) minimizes the average daily nutritional mismatch of the generated menu, whose elements are specified in constraints (5)–(13).

Constraints (5), (6), (7) and (8) limit the total daily energy input, a number of kilocalories contributed by each group of macronutrients each day, energy contribution of different meal schedules, and energy contribution of saturated and unsaturated fatty acids, respectively. Constraints (9) and (10) ensure that the requirements of vitamins and minerals in this study were satisfied according to the recommended and tolerable levels of intake, as specified by specialized organizations. In addition, there are elements that must be provided, although they are not considered as nutrients. Thus, constraint (11) controls the daily consumption of dietary fiber. Constraints (12) and (13) enable the proper daily and weekly intake of different food groups as suggested by experts. Constraints (14), (15), (16) and (17) specify the appropriate menus. Thus, constraint (14) requires that all dishes at different meal times on different days have an assigned food. Constraints (15) and (16) ensure that no main dish is served during two consecutive lunches or two consecutive dinners, respectively. Constraint (17) limits the size of portions that can be assigned.

Finally, constraints (18) and (19) define the types of variables in the model. The first variables were the assigned portion amount and mismatch levels, which must be larger than or equal to zero. The second set includes binary variables associated with the decision regarding whether to consider food under the established conditions. Then, the resulting model is a mixed integer linear programming problem.

## 3. Solution strategies

Unlike optimization problems with only one objective function, in the multiobjective case, a set of nondominated (efficient) solutions is sought after instead of an optimal solution. For example, if a multiobjective model includes several minimization objectives *Z_i_*(*x*), then a solution *y* dominates solution *x* if *Z_i_*(*y*) ≤ *Z_i_*(*x*) for every objective *i*, and there is at least one objective *i* such that *Z_i_* (*y*) < *Z_i_*(*x*). The set of solutions that are not dominated by another solution in the objective space is known as the Pareto border [9]. The model for MO-NMPP-CHO is solved using three different methods. The constraint method [20] is implemented using the General Algebraic Modeling System [21]; two multiobjective evolutionary algorithms (MOEA) are implemented in C++: Nondominated Sorting Genetic Algorithm II [22], which is also known as NSGA-II, and Strength Pareto Evolutionary Algorithm 2 [23], which is also known as SPEA2. To complete the numerical study, a set of test instances associated with boys and girls aged 4-18 years was created.

### 3.1 An approach for the MO-NMPP-CHO based on the constraint method

The purpose of the constraint method is to transform a multiobjective problem into several mono-objective problems to optimize one objective function, whereas those that become part of the constraints are limited by values ε. For example, let us consider the multiobjective model specified by equations (20) and (21), where objective *Z* is a vector of *p* functions *Z_i_*(*i* = 1, …, *p*), and *F_d_* is the feasible region. The constraint method generates several mono-objective models as illustrated in equations (22) – (24).

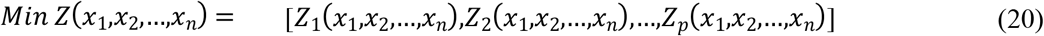

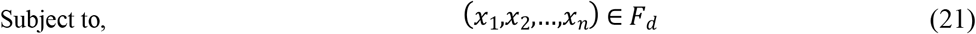

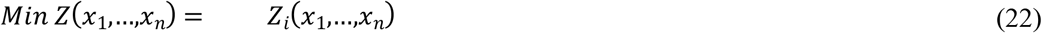

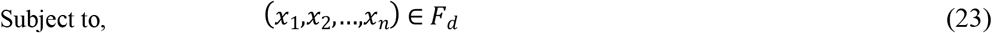

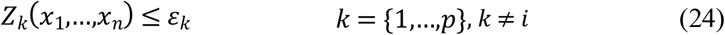

Our model includes *p* = 4 objective functions, and *F_d_* is specified by constraints (5)-(19). The described process is applied to each of the four objectives. The basic issue is to determine the appropriate values of ε*_i_* (*i* = 1, …, 4). Thus, a separate problem, as illustrated in equations (25) – (26), is solved for each objective function *Z_i_*, and the optimal solution 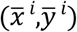 is used to specify the vector 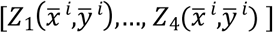. Then, the range of values 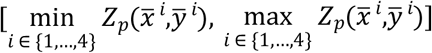 for each *ε_p_* (*p* = 1, …, 4) is divided into *t* parts to determine (*t*+1) values for *ε_p_*.

In our case, *t* = 2 generates 3 different values for *ε_p_*.

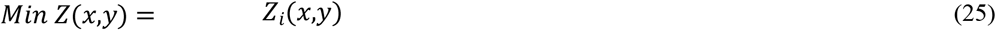

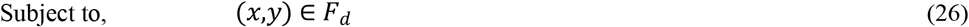

To complete the constraint method, for each objective *Z_i_* (*i* = 1, …, 4), the mono-objective model in equations (22) – (24) is solved for each combination of different values of *ε_k_*, *k* ϵ {1, …, 4}, where *k* ≠ *i* in their sets of values (i.e., 27 different problems are solved for each *i*). The models generated for different *ε* combinations are solved using the GAMS/CPLEX solver with the Branch-and-Cut algorithm. After the solutions for all of the models generated by the combination of *ε* values are found, the nondominance in the objective space is used over all solutions, which generates the Pareto border approximation.

### 3.2 Two evolutionary approaches for the MO-NMPP-CHO

In an evolutionary approach, a full population of solutions is modified during the process. Among these methods, a subclassification known as evolutionary algorithms presents multiple advantages to address multiobjective problems [24]. In fact, evolutionary algorithms are characterized by imitating the evolutionary process of the species regarding the survival of the fittest, i.e., a population of individuals (solutions to the problem) is modified after several generations through the application of parent-selection rules, crossover strategies and mutation strategies. Thus, the following series of elements must be introduced to proceed.

- Encoding the solution: definition of the coded representation (or chromosome) of individuals in the population in both the objective space and the decision space
- Fitness assignment: definition of a strategy to assign a value to each individual to motivate its aptitude to be part of the next generation
- Mating selection: definition of the strategy to select individuals to be parents of new solutions
- Environmental selection: definition of the strategy to decide the members of the current population that will be included in the population of the next generation
- Reproduction strategy: definition of the mutation and crossover operators to generate the next generation with the probability of applying each operator
- Initialization of population: Definition of the population size and strategy to create the initial population
- Stop criterion: definition of a criterion that enables the algorithm to stop after fulfilling a condition

We consider two evolutionary algorithms NSGA-II and SPEA2 to address the MO-NMPP-CHO focusing on childhood obesity. First, we defined the identical operators and strategies to implement both methods; then, we specified the different operators and particular strategies in each method.

<u>*Solution encoding*</u>: A solution in the decision space is represented using two rows and *T*columns, where *T* is the number of days multiplied by the number of dishes that should be served per day. Figure 1 illustrates the attributes of each row and column to encode the solution in the decision space. The first row includes the number of food portions to be served, and the second row includes an identifier of the food to be served. The position of each column considers different characteristics (e.g., the meal time to which it belongs and the dish in the meal). The representation of an individual in the objective space corresponds to a vector whose size is equal to the number of objectives (4 in this case).

**Fig 1.**
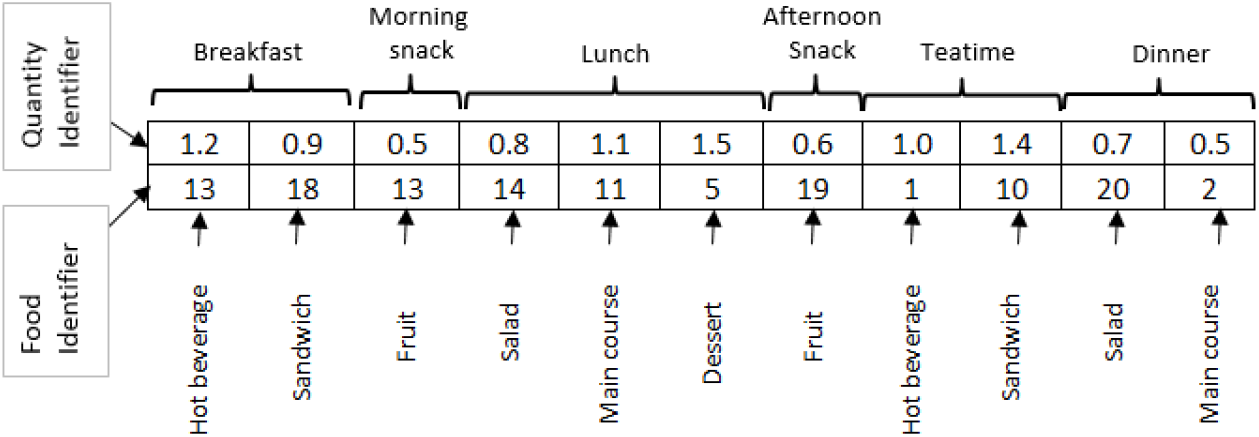
Representation of an individual in the decision space, planning for one day.

<u>*Reproduction strategy*</u>: For both metaheuristic methods, we used a crossover operator with *M*crossing points [19], where *M* is equal to the number of days to be planned. The mutation operator modifies the number of points equal to the number of days to be planned thus, both food and amount of food portion are randomly reallocated. Two different children are generated when the crossover operator is applied, and one child is randomly selected. A strategy of crossover-OR-mutation is used so that at least one operator (crossover or mutation) is applied during the crossbreeding application [25].

<u>*Initial population*</u>: To generate the initial population and ensure diversity within the objective space, individuals are randomly created. However, they become members of the population if they are not clones of any of the existing individuals in the initial population.

<u>*Stop criterion*</u>: The termination condition for both metaheuristic methods is the fulfillment of *G*_max_ generations or a maximum running time of 1,800 seconds.

#### 3.2.1 NSGA-II method for the MO-NMPP-CHO

To implement NSGA-II, the following elements must be specified. First, the fitness allocation is based on the dominance depth criterion that generates several layers in the population; that is, a population of individuals creates a better-quality layer (i.e., ranking 1), which includes individuals who are not dominated by others. The second layer with ranking 2, includes the individuals not dominated by others in the remaining population. The same principle applies to create other layers of higher ranking. The second element that specifies fitness is the density estimator, which is called the crowding distance and consists of estimating the perimeter of the cuboid formed by the neighbors closest to the individual in the objective space as illustrated in Figure 2 for a bi-objective maximization problem. Thus, the operator of crowding comparison specifies that an individual dominates another if it has a better ranking or equal ranking with a greater crowding distance.

**Figure 2:**
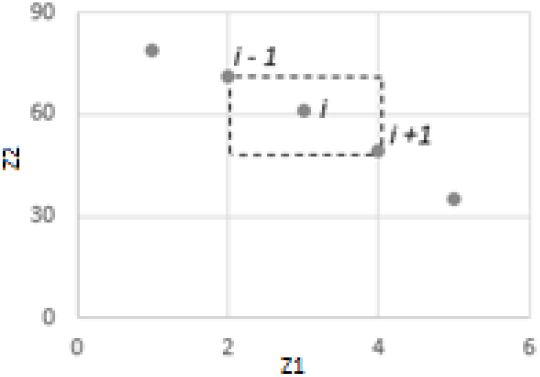
Example of crowding distance, where each point is a nondominate solution.

The parents to create the new population through the application of genetic operators are selected from the current population through a binary tournament using the crowding comparison operator to specify the fittest. Environmental selection is performed by adding the best layers from the current population to the new population until it reaches its size. If the population size cannot be achieved exactly, then individuals with better density indicators are added from the last candidate layer for inclusion until the size of the new population is attained. At the end of the procedure, the individuals with ranking 1 correspond to nondominate individuals and form the Pareto border approximation. We include the strategy of eliminating the overlapping solutions in the objective space after creating the new population as in [26]. Hence, in the worst case, *N* individuals are present instead of clones to continue the process because the initial population does not contain clones. To tune the parameters of NSGA-II a parameterization procedure was conducted using a 2^k^ factorial design [27] with a confidence level of 95% that resulted in the following parameters: population size, 300; the maximum number of generations, 500; and the probability of applying the crossover operator instead of the mutation operator, 95%.

#### 3.2.2 SPEA 2 method for the MO-NMPP-CHO

SPEA2 can be characterized as follows. Unlike NSGA-II, SPEA2 ensures elitism through an external file in addition to the main population of individuals. The size of the external file remains fixed because of the truncation operator; hence, when the sample exceeds the permitted size for the external file, individuals with a smaller distance to another individual in the objective space are iteratively eliminated until the sample has the permitted size, thereby avoiding the elimination of boundary solutions. The density estimator corresponds to the inverse of the Euclidean distance in the objective space between the individual and the *k*-th closest individual, where *k* is equal to the integer part of the square root of the sum of the size of the main population and the size of the external file. To obtain the fitness of an individual, it is necessary to calculate the strength value for each individual. Then, the fitness of an individual is equal to the sum of its raw fitness that corresponds to the sum of the strength value of the individuals who dominates it and its density estimator.

The environmental selection process to generate the external file of the next generation was applied by copying the nondominate individuals of the current main population and the external file into the external file of the next generation. The environmental selection process to generate the main population of the next generation was performed by applying genetic operators to the parents selected from the external file of the next generation. The parents were selected through a binary tournament using their fitness. When the stopping criterion was fulfilled, the individuals in the external file with fitness value less than one, corresponded to the nondominate individuals who formed the Pareto border approximation.

The parameterization For SPEA2 was performed using a 2^k^ factorial design with a 95% confidence level, which resulted in the following parameters: size of the main population, 300; maximum number of generations, 500; probability of applying the crossover operator instead of the mutation operator, 80%; and external file size: 50% of the main population size.

## 4. Results

The proposed mathematical programming model was solved with the three methods described in the previous section. For the constraint method, each problem instance was solved using GAMS software, version 24.3.3 with IBM ILOG CPLEX *Optimization Studio* solver, version 12.06.1 [21]. The metaheuristics were implemented in C++. All implementations were solved using a computer with an i7 Intel Core processor, 2.40 GHz and 8 GB of RAM.

### 4.1 Test instances

Because no instance was available in the literature to complete our numerical experimentation, a group of six instances with different age ranges under study was specified under the supervision of health professionals who have experience with the real situation. These instances were associated with children diagnosed with obesity across three age ranges: 4-8 years, 9-13 years and 14-18 years. Each designed instance differs in the amount of recommended daily energy, suggested dietary fiber, and recommended amounts for some or all of the micronutrients considered (see Table 2). The notations in Table 2 that characterize the instances include three elements: The first number indicates the lower limit of the age range; the letter indicates gender (“a”: girl; “o”: boy); and the second number indicates the upper limit of the age range, for example, 4o8 indicates a boy aged 4-8 years.

**Table 2:** Test Instances

|  |  | 4o8 | 4a8 | 9o13 | 9a13 | 14o18 | 14a18 |
| --- | --- | --- | --- | --- | --- | --- | --- |
| Energy [kJ/day] |  | 5857.6 | 5020.8 | 7531.2 | 6694.4 | 9204.8 | 7531.2 |
| Vitamin A [$\mu\text{g}/\text{day}$] | | 400/1300 | 400/1300 | 600/1700 | 600/1700 | 900/2800 | 700/2800 |
| Vitamin B1 [mg/day] |  | 0.6/*ND* | 0.6/*ND* | 0.9/*ND* | 0.9/*ND* | 1.2/*ND* | 1/*ND* |
| Vitamin B2 [mg/day] |  | 0.6/*ND* | 0.6/*ND* | 0.9/*ND* | 0.9/*ND* | 1.3/*ND* | 1/*ND* |
| Vitamin B3 [mg/day] |  | 8/15 | 8/15 | 12/20 | 12/20 | 16/30 | 14/30 |
| Vitamin B6 [mg/day] |  | 0.6/40 | 0.6/40 | 1/60 | 1/60 | 1.3/80 | 1.2/80 |
| Vitamin B9 [$\mu\text{g}/\text{day}$] | | 200/400 | 200/400 | 300/600 | 300/600 | 400/800 | 400/800 |
| Vitamin B12 [$\mu\text{g}/\text{day}$] | | 1.2/*ND* | 1.2/*ND* | 1.8/*ND* | 1.8/*ND* | 2.4/*ND* | 2.4/*ND* |
| Vitamin C [mg/day] |  | 25/650 | 25/650 | 45/1200 | 45/1200 | 75/1800 | 65/1800 |
| Vitamin E [mg/day] |  | 7/140 | 7/140 | 11/220 | 11/220 | 15/260 | 15/260 |
| Calcium [mg/day] |  | 1000/2500 | 1000/2500 | 1300/3000 | 1300/3000 | 1300/3000 | 1300/3000 |
| Copper [mg/day] |  | 0.44/3 | 0.44/3 | 0.7/5 | 0.7/5 | 0.89/11 | 0.89/8 |
| Iron [mg/day] |  | 8/11 | 8/11 | 8/11 | 8/11 | 11/20 | 15/20 |
| Magnesium [mg/day] |  | 150/250 | 150/250 | 300/400 | 300/400 | 300/400 | 300/400 |
| Phosphorus [mg/day] |  | 500/3000 | 500/3000 | 1250/4000 | 1250/4000 | 1250/4000 | 1250/4000 |
| Potassium [mg/day] |  | 2457/4500 | 2106/4500 | 3159/4500 | 2808/4500 | 3510/4700 | 3510/4700 |
| Selenium [$\mu\text{g}/\text{day}$] | | 15/40 | 15/40 | 15/40 | 15/40 | 40/55 | 40/55 |
| Sodium [mg/day] |  | 500/1400 | 500/1200 | 500/1800 | 500/1600 | 500/2000 | 500/1800 |
| Zinc [mg/day] |  | 5/12 | 5/12 | 8/23 | 8/23 | 9/24 | 11/24 |
| Dietary fiber [g/day] |  | 12/20 | 12/20 | 15/30 | 15/30 | 20/40 | 20/40 |

Table 3 indicates the parameters that do not depend on certain instances and are the valid nutritional recommendations for the 4-18 years. Thus, Table 3 includes recommendations for the proportion of energy contributed by each macronutrient, the proportion of energy contributed by each mealtime, the proportion of energy contributed by different fatty acids, and minimum/maximum allowed food portion. Table 3 also indicates the recommendations for daily and weekly food group consumption. Furthermore, a seven-day planning period was considered for all instances because the proposed model independently considers the weekly planning period.

**Table 3.** Recommendations for parameters: macronutrients, mealtimes, fatty acids, portion sizes, and food groups (daily and weekly)

| Parameter | Value | Food groups | $RGDI_g/RGDS_g$ | $RGSI_g/RGSS_g$ |
| --- | --- | --- | --- | --- |
| $PEMI_1/PEMS_1$ | 0.45/0.65 | Vegetable | 2/8 | 7/56 |
| $PEMI_2/PEMS_2$ | 0.10/0.30 | Fruit | 2/8 | 7/56 |
| $PEMI_3/PEMS_3$ | 0.20/0.35 | Dairy products | 2/8 | 7/56 |
| $ECl_1/ECS_1$ | 0.1/0.15 | Fish | 0/1 | 1/3 |
| $ECl_2/ECS_2$ | 0.05/0.1 | Red meat | 0/1 | 1/3 |
| $ECl_3/ECS_3$ | 0.3/0.4 | Poultry | 0/1 | 1/3 |
| $ECl_4/ECS_4$ | 0.05/0.1 | Egg | 0/1 | 1/3 |
| $ECl_5/ECS_5$ | 0.1/0.15 | Noodles | 0/1 | 2/5 |
| $ECl_6/ECS_6$ | 0.2/0.3 | Rice | 0/1 | 2/5 |
| $RL_1$ | 0.1 | Potatoes | 0/1 | 2/5 |
| $RL_2$ | 0.1 | Legume | 0/1 | 1/3 |
| $RL_4$ | 0.1 | | | |
| $RL_5$ | 0.012 | | | |
| $LIP/LSP$ | 0.5/1.5 | | | |

### 4.2 Numerical results

The methods were analyzed using the indicators proposed by Talbi [19], where || · || indicates the Euclidian distance in the objective space, and | · | indicates the cardinality of a set. To analyze the diversity within population *A* generated with a particular method, the Extent indicator *I_ex_*(*A*) was used (27), where *n* is the number of objective functions, *Z_i_*(*u*) is the value of the *i*-th objective function, and *Z*(*u*) is the vector of the objective functions of individual *u*.

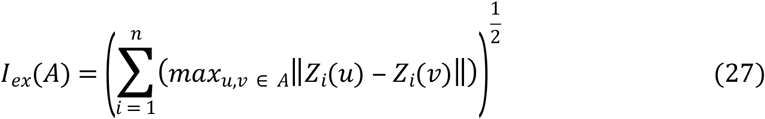

To measure the improvement obtained with heuristic methods, the generational distance *I_GD_*(*A,R*) that measures the distance between the final population *A* and the initial population *R* was used (28). The method with the best performance in terms of this indicator achieves the greatest distance between its initial and final populations.

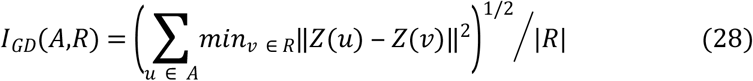

The Contribution indicator *Cont*(*PF*_1_/*PF*_2_), is used to measure the contribution of the nondominate solutions of two methods using approximations of the Pareto fronts *PF*_1_ and *PF*_2_. To combine the solutions from these methods, *PF* denotes the intersection of sets *PF*_1_ and *PF*_2_, *PF*\* includes the nondominate solutions in *PF*_1_ ∪ *PF*_2_, *W_1_* is the set of solutions of *PF*_1_ that dominates a solution in *PF*_2_, and *N_1_* corresponds to the set of solutions of *PF*_1_ that do not interact with the solutions of *PF*_2_ (i.e., solutions in *N_1_* that do not dominate any solution, are not dominated by any solution and are not clones of any solution of *PF*_2_). Finally, *Cont*(*PF*_1_/*PF*_2_) computes the proportion of nondominate solutions that *PF*_1_ gives to *PF*\*, *Cont*(*PF*_2_/*PF*_1_) computes the proportion of nondominated solutions that *PF*_2_ gives to *PF*\*, and *Cont*(*PF*_1_/*PF*_2_) + *Cont*(*PF*_2_/*PF*_1_) = 1. For example, if *Cont*(*PF*_1_/*PF*_2_) is greater than 0.5, then *Cont*(*PF*_2_/*PF*_1_) is less than 0.5; the method that generates *PF*_1_ is better than the method that generates *PF*_2_ in terms of convergence to the Pareto frontier.

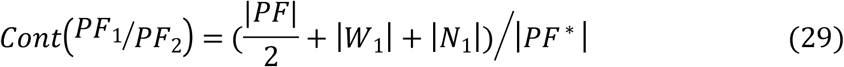

The evolutionary techniques required significantly less time to solve the proposed problem and found more solutions than the constraint method that uses the GAMS/CPLEX solver with Branch-and-Cut algorithm. Table 4 displays the number of solutions of the Pareto border approximation found with each method and the required execution time. Because the metaheuristics are stochastic processes, the average value of 10 executions is shown; for the constraint method, GAMS uses a deterministic Branch-and-Cut algorithm, so only a single execution is performed. Therefore, even for a small problem, the constraint method likely requires intensive calculations. Thus, this solution time prohibits a nutritionist working in the public health sector from attending to many patients during the day. Nevertheless, the population of solutions found using the constraint method indicates better behavior in terms of diversity in the objective space.

**Table 4:**
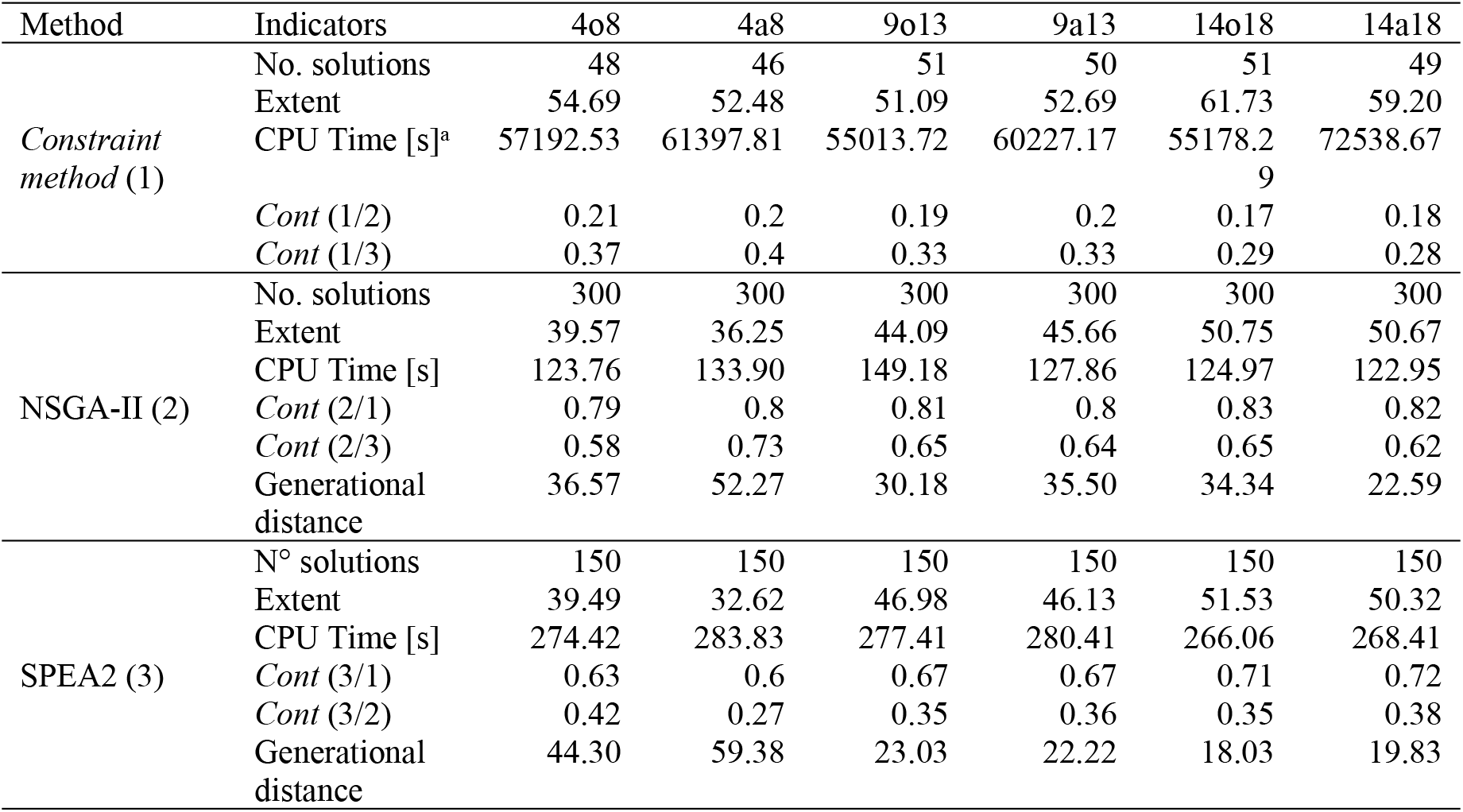
Computational performance of methods.

Contribution indicators were used to measure the proportion of individuals of a Pareto front approximation built by combining two populations. The contribution of the constraint method was less than 40%, although in all studied cases, the individuals of the Pareto front approximation generated using this method still belonged to that built from its combination with the population generated via the NSGA-II and SPEA2 methods. Thus, the evolutionary techniques presented better solutions to the Pareto front than the constraint method. The comparison of the performance of NSGA-II and SPEA2 did not differ in terms of average diversity indicator or average generational distance. However, even if NSGA-II requires only half the running time of SPEA2, the average difference was not greater than a few minutes. In terms of the contribution of the solutions to the approximation of the Pareto front created by combining the fronts of both methods, NSGA-II provided better results; however, this improvement might be affected by the population size.

The produced menus generally show that all methods repeat certain types of food with a strong relationship between price and nutritional benefit (e.g., skim milk, natural yogurt, and lentils with rice). Table 5 displays menus designed via the three methods for one day for a 4- to 8-year-old children. Each menu corresponds to an efficient solution. Notes that the evolutionary methods select many of the foods selected by the exact method.

**Table 5:** Menus created using the constraint method, NSGA-II and SPEA2.

| Mealtime | Dish | Method |  |  |  |  |  |
| --- | --- | --- | --- | --- | --- | --- | --- |
|  |  | Constraint method |  | NSGA-II |  | SPEA2 |  |
|  |  | Food | Portion | Food | Portion | Food | Portion |
| Breakfast | Hot beverage | Skim milk | 0.5 | Quaker oats with milk | 1.2 | Skim milk. | 0.6 |
|  | Sandwich | Whole wheat bread with ham | 0.5 | Whole wheat bread with honey | 0.9 | Whole wheat bread with margarine | 0.5 |
| Morning snack | Fruit | Strawberries | 0.6 | Cantaloupe | 1.0 | Cherries | 0.6 |
| Lunch | Salad | Lettuce | 1.4 | Broccoli | 1.0 | Celery salad | 1.4 |
|  | Main course | Conger chowder | 0.5 | Potatoes with egg. | 0.5 | Garbanzos. | 0.5 |
|  | Dessert | Natural yogurt | 0.5 | Plum | 1.1 | Blueberries. | 1.4 |
| Afternoon snack | Fruit | Cherries | 0.5 | Raspberries | 1.1 | Cherries | 0.6 |
| Teatime | Hot beverage | Skim milk | 0.5 | Quaker oats with milk | 0.6 | Quaker oats with milk | 0.8 |
|  | Sandwich | Whole wheat bread with ham | 0.5 | Whole wheat bread with ham | 0.7 | Whole wheat bread with fresh cheese | 0.5 |
| Dinner | Salad | Lettuce | 1.0 | Green beans | 0.5 | Celery salad | 1.4 |
|  | Main course | Lentils with rice | 0.5 | Hake with Potato | 1.2 | Charquicán | 0.5 |
| Objectives functions values |  |  |  |  |  |  |  |
| $Z_1$ [USD] | | 4.76 | | 2.53 | | 3.78 | |
| $Z_2$ [mg] | | 96.1 | | 143.5 | | 105.5 | |
| $Z_3$ [u] | | 60.8 | | 72.2 | | 70.2 | |
| $Z_4$ [%] | | 0 | | 12 | | 7 | |

## 5. Conclusions

The multiobjective approach proposed in this paper generates menus that minimize the consumption of substances that are particularly harmful to obese children. Also, it minimizes the nutritional mismatch and cost of planning to avoid limited access to healthy diets because of economic issues, while complying with the nutritional recommendations of specialized organizations. The proposed model for MO-NMPP-CHO with the created instances was solved with a deterministic method and two metaheuristic methods.

Although childhood obesity is a multifactor problem, the formation of healthy eating habits at an early age presents benefits over the long term. Thus, the multiobjective mathematical programming model for planning nutritional menus in this paper appears to be an appropriate way to minimize exposure to the major risk factors for the development of chronic diseases associated with childhood obesity, the total cost of nutritional planning, and nutritional mismatch.

Nevertheless, the numerical results indicate that solving this type of problem using exact methods is not appropriate to address real or complex instances because of their execution time. Positive results can be obtained using evolutionary techniques that require appropriate computational times. Although these techniques provide only an approximate analysis, public health professionals can use them as a guide to achieving personalized plans based on the requirements of each child.

## ACKNOWLEDGMENTS

The CONICYT-BASAL partially supported this study [grant number FB0816].

